# An epigenome-wide association study of child appetitive traits and DNA methylation

**DOI:** 10.1101/2023.07.17.549289

**Authors:** Holly A. Harris, Chloe Friedman, Anne P. Starling, Dana Dabelea, Susan L. Johnson, Bernard F. Fuemmeler, Dereje Jima, Susan K. Murphy, Cathrine Hoyo, Pauline W. Jansen, Janine F. Felix, Rosa Mulder

**Author notes:** **Address correspondence to:** Holly A. Harris, Erasmus University Rotterdam, Department of Psychology, Education & Child Studies, P.O. Box 1738, 3000 DR Rotterdam, The Netherlands. (H.A. Harris), (C. Friedman), (A. Starling), (D. Dabelea), (S.L. Johnson), (B. F. Fuemmeler), (D. Jima), (S. K. Murphy), (C. Hoyo), (P.W. Jansen), (J. F. Felix), (R. Mulder).

## Abstract

Childhood appetitive traits are consistently associated with obesity risk, and yet their etiology is poorly understood. Appetitive traits are complex phenotypes which are hypothesized to be influenced by both genetic and environmental factors, as well as their interactions. Early-life epigenetic processes, such as DNA methylation (DNAm), may be involved in the developmental programming of appetite regulation in childhood. In the current study, we meta-analyzed epigenome-wide association studies (EWASs) of cord blood DNAm and early-childhood appetitive traits. Data were from two independent cohorts: the Generation R Study (*n*=1,086, Rotterdam, the Netherlands) and the Healthy Start study (*n*=236, Colorado, USA). DNAm at autosomal methylation sites in cord blood was measured using the Illumina Infinium HumanMethylation450 BeadChip. Parents reported on their child’s food responsiveness, emotional undereating, satiety responsiveness and food fussiness using the Children’s Eating Behaviour Questionnaire at age 4-5 years. Multiple regression models were used to examine the association of DNAm (predictor) at the individual site- and regional-level (using DMRff) with each appetitive trait (outcome), adjusting for covariates. Bonferroni-correction was applied to adjust for multiple testing. There were no associations of DNAm and any appetitive trait at the individual site-level. However, at the regional level, we identified 45 associations of DNAm with food responsiveness, 7 associations of DNAm with emotional undereating, 13 associations of DNAm with satiety responsiveness, and 9 associations of DNAm with food fussiness. This study shows that DNAm in the newborn may partially explain variation in appetitive traits expressed in early childhood and provides preliminary support for early programming of child appetitive traits through DNAm. Investigating differential DNAm associated with appetitive traits could be an important first step in identifying biological pathways underlying the development of these behaviors.

## 1. Introduction

Appetitive traits (or ‘eating behaviors’) are associated with the development of obesity (Llewellyn & Fildes, 2017). Meta-analytic evidence from infant and child studies show robust associations between ‘food approach’ traits and higher obesity risk, while ‘food avoidant’ traits confer a lower obesity risk (Kininmonth et al., 2021). This may, in part, be explained through dietary behaviors. Food approach traits, for example food responsiveness, indicate a child’s appetitive avidity, and sensitivity to respond to environmental food cues. Food responsiveness is associated with a greater preference for energy-dense and nutrient-poor foods (Fildes et al., 2015), increased eating frequency and higher energy intake (Carnell & Wardle, 2007; Syrad et al., 2016). In contrast, food avoidant behaviors, such as food fussiness, emotional undereating and satiety responsiveness, may reflect a lower appetite and disinterest in eating. Children high in satiety responsiveness are sensitive to feelings of fullness, tend to eat slower and eat smaller portions (Carnell & Wardle, 2007; Syrad et al., 2016). Children with elevated food fussiness tend to consume a limited variety of foods (Taylor et al., 2015) and tend to dislike core food groups such as fruits and vegetables (Fildes et al., 2015). Finally, children high in emotional undereating eat less when negative moods are evoked (Blissett et al., 2019). While appetitive traits appear to bridge associations between nutrition, the food environment and weight, relatively little is known about the early developmental processes underlying appetitive traits.

Appetitive traits themselves are complex phenotypes that are influenced by genetic and environmental factors, as well as their interactions. Individual differences in appetitive traits emerge early in postnatal life (Llewellyn et al., 2011) and remain moderately stable into the early childhood years (Costa et al., 2022; Farrow & Blissett, 2012). Twin heritability studies suggest that genetic variants play a considerable role in explaining variation in some appetitive traits (Llewellyn & Wardle, 2015).

The prenatal environment has also been hypothesized to contribute to obesity-risk, partly through modulating appetite regulation systems linked to appetite trait expression (Boswell et al., 2018; Desai & Ross, 2020). Obesity risk factors during pregnancy (such as high fasting plasma glucose and pre-pregnancy overweight) have been shown to predict eating behaviors such as self-serving larger portions, faster eating rate and higher energy intake, as well as increased child body mass index (BMI), at 6 years (Fogel et al., 2020). Observational evidence from a longitudinal birth cohort showed that excessive gestational weight gain was associated with increased food responsiveness in children aged 1 year old (Costa et al., 2022). Another study showed that mothers who gained less weight than recommended during pregnancy rated their male (but not female) offspring lower in satiety responsiveness at approximately 4 years of age (Boone-Heinonen et al., 2019). Furthermore, ultra-processed food consumption during pregnancy has been also linked to lower satiety responsiveness in infants at 6-months of age (Cummings et al., 2022). The prenatal environment may be linked to children’s appetitive traits through biological processes that could modify the regulation or expression of genes during fetal development, such as epigenetic processes.

DNA methylation (DNAm) is an epigenetic mechanism whereby methyl groups are added to cytosines at cytosine-guanine dinucleotides (CpG sites) in the DNA, which may affect gene expression (Ruiz-Arenas et al., 2022). Exposures during fetal development, such as maternal smoking (Joubert et al., 2016), maternal BMI at the start of pregnancy (Sharp et al., 2017) and maternal diet (Küpers et al., 2022) have been associated with newborn DNAm. A limited number of candidate studies have linked DNAm to appetitive traits in childhood. For example, one study (*n*=317) showed that DNAm at the insulin-like growth factor-2 (*IGF2*) gene in cord blood was associated with satiety responsiveness in children aged 1-6 years old (Do et al., 2019). In a small study of 32 girls aged 5-6 years, lower levels of DNAm in the promoter of the *BDNF* (brain-derived neurotrophic factor) gene were found to be associated with lower satiety responsiveness (Gardner et al., 2015). While such studies implicate differential DNAm as a potentially relevant pathway underlying appetitive traits, so far interest has been in one or a few candidate genes, thereby precluding the possibility to uncover novel epigenetic pathways.

Epigenome-wide association studies (EWAS) of DNAm markers at birth that predict childhood appetitive traits have not yet been performed. Hypothesis-free testing of epigenome-wide DNAm signals may present new leads to piece together the puzzle of appetitive trait etiology. Thus, in the current study, we aimed to investigate associations of genome-wide DNAm in newborn cord blood with appetitive traits in early childhood. To maximize statistical power, we meta-analyzed EWASs for appetitive traits from two independent cohorts (total *n*=1,322).

## 2. Methods

### 2.1. Study design and participants

Two cohorts participating in the worldwide Pregnancy And Childhood Epigenetics Consortium (PACE) (Felix et al., 2018) with newborn umbilical cord blood DNAm and child appetitive traits measured before age 6 years were included in the current analysis: the Generation R Study (Generation R) and Healthy Start.

#### 2.1.1. Generation R

Generation R is a population-based cohort focused on health and development from fetal life onwards (Jaddoe et al., 2006). All pregnant women living in Rotterdam, the Netherlands, with an expected delivery date between April 2002 and January 2006 were invited to participate (*N*=9,778; participation rate: 61%). Cord blood DNAm data were available for 1,396 children, who were selected from the full population to be a homogeneous subgroup of European ancestry, with high completeness of follow-up. Of these children, 1,098 had information available on appetitive traits. Twelve sibling pairs were present in this set; one sibling of each pair was removed based on data availability or otherwise randomly, leaving a final sample size of *n*=1,086 children. The Generation R Study has been approved by the Medical Ethical Committee of Erasmus MC, University Medical Center Rotterdam. Written informed consent was obtained from parents of all children.

#### 2.1.2. Healthy Start

The Healthy Start study (Healthy Start) is an ongoing, pre-birth cohort study based in Colorado, USA (Starling et al., 2015). Pregnant women were recruited between 2009-2014 at the University of Colorado obstetrics clinics. Women were eligible for the Healthy Start study if they were ≥16 years, expecting a singleton birth, had a gestational age <24 weeks at enrolment, and had no serious chronic medical conditions or history of stillbirth. A total of 1,410 pregnancies were enrolled, and umbilical cord blood DNAm was analyzed in a subset of these (*n*=600), based on availability of cord blood, maternal blood, and urine samples during pregnancy. For this analysis of cord blood DNAm and child appetitive traits, further exclusions were as follows: 6 participants had discordance between predicted and reported sex, 180 had missing child appetitive traits data, 7 were siblings of other participants in the cohort, and 171 had a race/ethnicity other than non-Hispanic white, resulting in a final sample size of *n*=236. The Healthy Start study protocol was approved by the Colorado Multiple Institutional Review Board, and all women provided written informed consent before the first study visit.

### 2.2. DNA methylation measurement

Each cohort performed sample processing, quality control and normalization based on their own protocols. In Generation R, DNA was extracted from cord blood samples taken at birth. Five-hundred nanograms of DNA per sample underwent bisulfite conversion using the EZ-96 DNA Methylation kit (Shallow) (Zymo Research Corporation, Irvine, USA) and further processed with the Illumina Infinium HumanMethylation450 BeadChip (Illumina Inc., San Diego, USA). Preparation and normalization of the HumanMethylation450 BeadChip array data was performed following the CPACOR workflow using the software package R (Lehne et al., 2015; R Core Team, 2013). In detail, the .idat files were read using the minfi package (Aryee et al., 2014). Probes with a detection *p*-value>1e-16 were set to missing per array. Intensity values were stratified by autosomal and non-autosomal probes and quantile normalized for each of the six probe type categories separately: type II red or green, type I methylated red or green and type I unmethylated red or green. Beta values were computed as the ratio of the methylated to the methylated+unmethylated signal. Arrays with observed technical problems such as failed bisulfite conversion, hybridization or extension, as well as arrays with a mismatch between sex of the proband and sex determined by the chromosome X and Y probe intensities were removed from subsequent analyses. Lastly, only arrays with a call rate >95% per sample were processed further. Outlying values on the probes were excluded using the Tukey method, i.e. values<(25^th^ percentile–3*interquartile range) and values>(75^th^ percentile+3*interquartile range) were excluded (Tukey, 1977).

In Healthy Start, DNA was extracted from umbilical cord blood collected at birth using the QIAamp DNA Blood Mini Kit (Qiagen) and stored in buffy coats. Illumina Infinium HumanMethylation450 BeadChip (Illumina Inc., San Diego, USA) was used to measure epigenome-wide DNAm (Yang et al., 2017). Additional details of DNA extraction and bisulfite conversion have been described in detail previously (Starling et al., 2020). Quality control checks included assessment of DNA purity, integrity, and quantity. Samples were eligible for inclusion if they met the following criteria: 260/280 ratio >1.8 indicating DNA purity, DNA integrity score >7, and at least 500 ng of DNA available. Probes with high detection p-value (>0.05) (n=587) and low beadcount <4 (n=664) were removed. Additionally, samples with inconsistencies between reported and predicted sex were removed (n=6). Stratified quantile normalization was performed using the preprocessQuantile function in minfi (Touleimat & Tost, 2012). ComBat was used for batch correction (Johnson et al., 2007). Extreme methylation outliers were removed, as defined by a value more than three times the interquartile range below the 25th percentile or above the 75^th^ percentile (Hoaglin et al., 1986; Merid et al., 2020).

For both cohorts, only autosomal probes were analyzed and cross-reactive probes were excluded (Chen et al., 2013; Naeem et al., 2014), resulting in 415,786 probes in Generation R and 429,136 probes in Healthy Start. We included the 415,267 probes in the meta-analyses that were present in both cohorts. The R package FDb.InfiniumMethylation.hg19 (Triche, 2014) was used for probe annotation.

### 2.3. Measures

#### 2.3.1. Child appetitive traits

In each cohort, child appetitive traits were assessed by parent-report using the widely-implemented Children’s Eating Behaviour Questionnaire (CEBQ) (Wardle et al., 2001) at age 4 years in Generation R, and age 5 years in Healthy Start. The CEBQ has good psychometric properties (Wardle et al., 2001), and has demonstrated ecological validity in behavioural tests (Blissett et al., 2019; Carnell & Wardle, 2007). Four subscales that were available in both Generation R and Healthy Start were included in the EWAS meta-analyses: food responsiveness (5 items, e.g. “*My child is always asking for food*”), emotional undereating (4 items, e.g. *“My child eats less when (s)he is upset”*), satiety responsiveness (5 items, e.g. *“My child gets full before his/her meal is finished”*) and food fussiness (6 items, e.g. *“My child is difficult to please with meals”*). Items were assessed on a 5-point Likert scale from “1” (never) to “5” (always) and summed to produce a subscale sum score. The subscales showed acceptable to good internal reliability in Generation R (α=0.74-0.91) and Healthy Start (α=0.70-0.93). Where normally distributed, sum scores were standardized (z-scores) for comparison between the models. The food responsiveness subscale was skewed in both cohorts and, therefore, square-root transformed to approach normality, and all scales were z-score transformed.

#### 2.3.2. Covariates

In Generation R, maternal age at delivery was calculated based on maternal age assessed upon enrolment, gestational age at enrolment and gestational age at birth. Maternal smoking during pregnancy was assessed with three questionnaires in early (<18 weeks), mid-(18-25 weeks), and late (>25 weeks) pregnancy and categorized into 0=never smoked during pregnancy/1=quit when pregnancy was known/2=sustained smoking during pregnancy. Maternal education was used as an indicator of socio-economic status (SES) and was obtained via questionnaire at enrolment. The information was dichotomized (0=did not complete university studies/1=completed university studies). Maternal pre-pregnancy body mass index (BMI; kg/m^2^) was computed based on pre-pregnancy weight, which was collected at enrolment using a questionnaire, and height. Self-reported pre-pregnancy weight correlated highly (*r*=0.96) with measured weight at enrolment around 13 weeks of pregnancy (Tielemans et al., 2015). Child sex and birth weight were obtained from midwife and hospital registries. Sample plate was included in the models to correct for batch effects. White blood cell proportions (CD4+ T-lymphocytes, CD8+ T-lymphocytes, natural killer cells, B-lymphocytes, monocytes, granulocytes, and nucleated red blood cells) were estimated with a cord blood-specific reference panel (Gervin et al., 2019). Gestational age at birth was determined using fetal ultrasound examinations or last menstrual period (LMP) in case of a regular menstrual cycle and a certain first day of LMP (Jaddoe et al., 2006).

In the Healthy Start study, child sex, birthweight, and gestational age at birth were obtained from medical records. Maternal age, ethnicity, and education were self-reported at time of enrolment. Education was dichotomized (0= high school education or less/ 1= more than high school education. Maternal smoking status was self-reported at three study visits: twice during pregnancy and once at delivery, and was operationalized as a 3-level categorical variable (0=never smoked/quit early in pregnancy, 1=quit smoking early in pregnancy, 2=sustained smoking throughout pregnancy). Maternal pre-pregnancy BMI was calculated using height measured at enrolment and pre-pregnancy weight obtained from the medical record. Cell deconvolution to estimate the relative proportions of the 7 cell types (B cells, CD4 T cells, CD8 T cells, granulocytes, monocytes, NK cells, and nucleated red blood cells) was performed using the R package FlowSorted.CordBloodCombined.450k (R Core Team, 2013) with a combined reference dataset for cord blood (Gervin et al., 2019).

### 2.4. Statistical Analyses

#### 2.4.1. Site- and regional-level epigenome-wide meta-analyses

Each cohort first ran a cohort-specific EWAS according to a predefined analysis plan. Missing data points on covariates were imputed (% missing on variables ranged from 0-9.8% in Generation R and 0-16.9% in Healthy Start) using the *mice* package (Buuren & Groothuis-Oudshoorn, 2010) in R. To enhance imputation, additional variables were used for the imputation only, including household income, maternal and paternal BMI at enrolment, maternal daily caloric intake during pregnancy and maternal alcohol intake during pregnancy in Generation R, and gestational weight gain and maternal daily caloric intake in Healthy Start. We performed a maximum of 10 iterations to create 10 imputed datasets. Regression analyses were performed using pooled statistics. To study the associations of DNAm with appetitive traits, we planned three regression models per appetitive trait: a basic model (Model 1), a fully adjusted model (Model 2), and an exploratory model including potential mediators (Model 3). In each model, DNAm was modelled as the predictor and appetitive trait as the outcome. Model 1 adjusted for child sex and age at appetitive trait assessment, batch and estimated white blood cell proportions. Significant hits (Bonferroni corrected, see below) were followed up by running Model 2, which additionally adjusted for potential confounders maternal age at delivery, smoking during pregnancy, education, and pre-pregnancy BMI. If Model 2 produced an association, Model 3 was run, which additionally adjusted for the potential mediators gestational age at birth and birth weight. With these models, site-level epigenome-wide analyses (EWASs) were performed separately in each cohort. Associations between DNAm at each individual site and standardized child appetitive trait score were estimated in R using robust linear regressions. Within each model, a Bonferroni-correction was applied to account for the number of probes (*p<*1.20×10^-7^). Quality control procedures on the EWAS results were run using the QCEWAS R package (Van der Most et al., 2017). Fixed effects inverse variance weighted meta-analyses of the cohort-specific results were performed using METAL (Willer et al., 2010). Shadow meta-analyses were independently performed a different investigator on the team and results were confirmed. For the regional-level EWASs, summary statistics from each cohort site-level EWASs were re-analyzed while taking the cohort specific-intercorrelational DNAm structure across sites into account, using the meta-analysis extension of the DMRff package in R (Suderman et al., 2018). This package identifies differentially methylated regions by taking into account dependencies between sites as well as site-level uncertainty in EWAS estimates. We searched within a standard 500bp site-window, applying a Bonferroni corrected *p*<0.05 based on the number of regions.

#### 2.4.2. Genetic enrichment

Significant findings stemming from site- or regional-level analyses were further examined in two ways. Genetic influences on DNAm were examined by i) a look-up of known associations of DNAm with genetic variants elsewhere on the genome, e.g. methylation quantitative trait loci (mQTLs) in *cis* (within a ±1 Mb window) or in *trans* (outside of this window, potentially on a different chromosome) as identified in cord blood of a pediatric population (Gaunt et al., 2016) and in cord or whole blood in a larger study on 36 populations of all ages (Min et al., 2020); and ii) a look-up of estimated additive genetic influences, and shared and unique environmental influences on DNAm, as based on twin heritability analyses (Hannon et al., 2018). To understand if there was enrichment of genetic patterns in associated versus non-associated CpGs, two-sided T-tests were performed between these two groups for each look-up and differences of *p*<0.05 are reported.

#### 2.4.3. Enrichment for eQTMs

We examined associated expression patterns by checking if significant findings have been identified as expression quantitative trait methylation (eQTMs) in *cis* in peripheral blood of 832 children aged between 6 and 11 years (Ruiz-Arenas et al., 2022). To understand if there was enrichment of gene expression patterns in associated versus non-associated CpGs, two-sided T-tests were performed between these two groups and differences of *p*<0.05 are reported.

#### 2.4.4. Enrichment for regulatory elements

We explored enrichment of tissue- or cell-type specific regulatory elements. This analysis was performed with eFORGe v2.0, using data from Consolidated Roadmap Epigenomics, ENCODE, and Blueprint to test for DNAse I hypersensitive regions, chromatin states, and histone marks, using default settings (1 kb window, 1000 background repetitions) (Breeze et al., 2019). Findings of FDR adjusted *q*<0.05 are reported.

#### 2.4.5. Functional enrichment

To examine functional enrichment of genes associated with sites significant in the site- or regional-level meta-analyses, we performed appetitive trait-specific Gene Ontology pathways analyses using the GOfuncR package in R (Grote & Grote, 2018). Pathways with a family-wise error rate (FWER) corrected *p*<0.05 based on random permutations of the gene-associated variables were considered to be enriched.

## 3. Results

The sample characteristics of Generation R and Healthy Start are described in **Table 1**. Both cohorts were comparable in their characteristics and appetitive traits. However, relative to Generation R, the Healthy Start children were slightly older (4.0 years *vs.* 4.9 years, respectively). In Generation R, mothers were slightly older (32.5 years *vs.* 30.4 years, respectively) and had a lower pre-pregnancy BMI (23.2 kg/m^2^ *vs.* 24.8 kg/m^2^, respectively) compared to Healthy Start mothers.

**Table 1.** Sample characteristics.

|  | *Generation R*<br>(*n*=1086) | *Healthy Start*<br>(*n*=236) |
| --- | --- | --- |
| **Child** | *n* (%) or mean ± SD | |
| Sex (% boys) | 531 (49) | 125 (53) |
| Birthweight (g) | 3568 ± 490 | 3333 ± 442 |
| Gestational age, weeks | 40.2 ± 1.4 | 39.6 ± 1.2 |
| Age at appetitive traits assessment, months | 48.5 ± 1.0 | 58.9 ± 9.9 |
| Appetitive traits<sup>a</sup> |  |  |
| Food responsiveness (scale 5 to 25) | 8.9 ± 3.1 | 11.1 ± 3.1 |
| Emotional undereating (scale 5 to 20) | 11.1 ± 3.4 | 10.7 ± 3.4 |
| Satiety responsiveness (scale 5 to 25) | 15.2 ± 3.2 | 15.9 ± 2.8 |
| Food fussiness (scale 5 to 30) | 17.8 ± 4.9 | 17.7 ± 5.1 |
| **Mother** | *n* (%) or mean ± SD | |
| Age (years)<sup>b</sup> | 32.5 ± 4.0 | 30.4 ± 4.8 |
| Pre-pregnancy BMI, (kg/m<sup>2</sup>) | 23.2 ± 3.8 | 24.8 ± 5.3 |
| Smoking |  |  |
| Never | 874 (80.5) | 171 (72.5) |
| Quit when pregnancy was known | 92 (8.5) | 10 (4.2) |
| Continued during pregnancy | 118 (10.9) | 15 (6.4) |
| Educational level<sup>c</sup> |  |  |
| High | 451 (41.5) | 208 (88.1) |

BMI: Body Mass Index; <sup>a</sup>Sum scores from the Children's Eating Behaviour Questionnaire (Wardle et al., 2001); <sup>b</sup>In Generation R, maternal age is assessed at delivery. In Healthy Start, maternal age is measured at enrolment; <sup>c</sup>In Generation R, mothers were coded as having a 'high' education if they completed university; in Healthy Start, mothers were coded as 'high' education if they completed more than a high school education; % missing on variables ranged from 0-9.8% in Generation R and 0-16.9% in Healthy Start

### 3.1. Site-level and regional-level epigenome-wide meta-analyses

We did not find associations of DNAm at individual CpG-sites and any of the appetitive traits in Model 1. Therefore, no further analyses were conducted relating to DNAm of individual CpG-sites. However, we found widespread associations of differentially methylated *regions* across the genome with each of the appetitive traits in Model 1. Most regions in Model 1 were also detected as a region in Model 2 (91% detected; differences may occur if inter-CpG similarity in effect size changes between models) and the majority remained significant (94% of detected regions). The results for each model are detailed in **Supplemental Tables 1-4**; here we present the results of fully adjusted Model 2. DNAm at 45 regions was associated with food responsiveness (**Table 2**), at 7 regions with emotional undereating (**Table 3**), at 13 regions with satiety responsiveness (**Table 4**), and at 9 regions with food fussiness (**Table 5**). DNAm at several regions was associated, in a consistent direction, with multiple appetitive traits: DNAm at a region on chromosome 4, near *HS6ST1,* was associated with both emotional undereating and satiety responsiveness; DNAm at a region on chromosome 12 in *GLIPR1L2*, and on chromosome 4 in *SH3BP2* was associated both with satiety responsiveness and food fussiness. The additional adjustment for maternal age, smoking, education, and pre-pregnancy BMI in Model 2 resulted in small effect size changes between Model 1 and Model 2 across all appetitive traits (median change=1.0%; IQR=0.4-2.2%). In Model 3, which included additional adjustment for the potential mediators gestational age at birth and birth weight, 86% of regions significant in Model 2 were detected. Of these, 88% remained significant (31 regions for food responsiveness; 5 regions for emotional undereating; 12 regions for satiety responsiveness; 8 regions for food fussiness). Effect size changes from Model 2 to Model 3 were small (median change=2.4%; IQR=1.4-3.8%), suggesting limited or no mediation of effects via changes in birth weight and gestational age at birth.

**Table 2.** Associated regions for epigenome-wide association study of DNA methylation (of cord blood at birth) and food responsiveness at 4-5 years.

| Chromosome | Start (hg19) | End (hg19) | Nearest gene | *n* CpGs | Estimate | SE | *p* | *p*<br>Bonferroni adjusted |
| --- | --- | --- | --- | --- | --- | --- | --- | --- |
| chr13 | 31506376 | 31507139 | *TEX26/TEX26-AS1* | 9 | 18.81 | 2.00 | 6.16E-21 | 3.16E-15 |
| chr17 | 14206774 | 14207968 | *HS3ST3B1* | 13 | 26.92 | 3.36 | 1.09E-15 | 5.58E-10 |
| chr20 | 44440824 | 44441164 | *UBE2C* | 6 | 37.24 | 4.75 | 4.68E-15 | 2.40E-09 |
| chr19 | 49223814 | 49224454 | *RASIP1* | 6 | 24.60 | 3.21 | 1.92E-14 | 9.83E-09 |
| chr11 | 3819041 | 3819306 | *PGAP2* | 6 | 215.24 | 30.55 | 1.85E-12 | 9.50E-07 |
| chr13 | 113622633 | 113622750 | *MCF2L-AS1* | 6 | 26.74 | 3.49 | 1.74E-14 | 8.91E-09 |
| chr11 | 44332385 | 44332940 | *ALX4* | 20 | 58.52 | 8.91 | 5.21E-11 | 2.67E-05 |
| chr11 | 36422377 | 36422615 | *PRR5L* | 5 | 13.33 | 2.01 | 3.60E-11 | 1.84E-05 |
| chr11 | 843915 | 844536 | *TSPAN4* | 8 | 24.67 | 3.80 | 8.70E-11 | 4.45E-05 |
| chr7 | 94023822 | 94023869 | *COL1A2* | 3 | 187.16 | 28.88 | 9.13E-11 | 4.67E-05 |
| chr8 | 145651369 | 145651799 | *VPS28* | 3 | -62.99 | 9.68 | 7.63E-11 | 3.91E-05 |
| chr9 | 90589146 | 90589806 | *CDK20* | 5 | 106.99 | 16.97 | 2.86E-10 | 1.47E-04 |
| chr1 | 156307962 | 156308296 | *CCT3/TSACC* | 9 | 161.86 | 26.48 | 9.81E-10 | 5.02E-04 |
| chr7 | 32111062 | 32111068 | *PDE1C* | 3 | 77.98 | 12.62 | 6.47E-10 | 3.31E-04 |
| chr4 | 5021084 | 5021311 | *CYTL1* | 7 | -23.84 | 4.04 | 3.59E-09 | 1.84E-03 |
| chr6 | 149805995 | 149806339 | *ZC3H12D* | 7 | 22.41 | 3.65 | 7.99E-10 | 4.09E-04 |
| chr11 | 2919798 | 2920209 | *SLC22A18AS* | 8 | 35.84 | 6.13 | 4.95E-09 | 2.54E-03 |
| chr12 | 45270312 | 45270573 | *NELL2* | 6 | 136.55 | 22.87 | 2.37E-09 | 1.22E-03 |
| chr15 | 90208810 | 90209326 | *PLIN1* | 5 | 104.26 | 17.56 | 2.91E-09 | 1.49E-03 |
| chr3 | 48471300 | 48471771 | *PLXNB1* | 5 | 52.86 | 8.98 | 3.90E-09 | 2.00E-03 |
| chr1 | 151118299 | 151119525 | *SEMA6C* | 13 | 93.74 | 16.17 | 6.73E-09 | 3.45E-03 |
| chr13 | 111039873 | 111040579 | *COL4A2* | 4 | 13.97 | 2.38 | 4.19E-09 | 2.15E-03 |
| chr22 | 18985500 | 18985691 | *DGCR5* | 3 | 12.75 | 2.20 | 6.50E-09 | 3.33E-03 |
| chr8 | 1765074 | 1766126 | *MIR596* | 13 | 27.14 | 4.67 | 6.23E-09 | 3.19E-03 |
| chr1 | 201476362 | 201476619 | *CSRP1* | 6 | 51.06 | 8.93 | 1.09E-08 | 5.59E-03 |
| chr11 | 567966 | 568206 | *MIR210HG* | 4 | 65.83 | 11.42 | 8.17E-09 | 4.19E-03 |
| chr11 | 111741914 | 111742291 | *ALG9* | 7 | 232.34 | 41.26 | 1.80E-08 | 9.20E-03 |
| chr17 | 76976010 | 76976357 | *LGALS3BP* | 7 | 41.90 | 7.27 | 8.31E-09 | 4.25E-03 |
| chr17 | 39890756 | 39891009 | *JUP* | 4 | 255.27 | 44.57 | 1.02E-08 | 5.23E-03 |
| chr7 | 79083054 | 79084166 | *MAGI2-AS3* | 17 | 36.17 | 6.41 | 1.67E-08 | 8.57E-03 |
| chr5 | 140018644 | 140018709 | *TMCO6* | 3 | 27.53 | 4.87 | 1.59E-08 | 8.12E-03 |
| chr5 | 140621375 | 140621754 | *PCDHB19P* | 3 | 26.72 | 4.81 | 2.75E-08 | 1.41E-02 |
| chr18 | 9708890 | 9709061 | *RAB31* | 3 | 376.43 | 67.81 | 2.83E-08 | 1.45E-02 |
| chr2 | 239140032 | 239140340 | *TARDBPP3/LINC02610* | 7 | 63.12 | 11.08 | 1.21E-08 | 6.18E-03 |
| chr7 | 35293080 | 35293892 | *TBX20* | 11 | 75.25 | 14.03 | 8.24E-08 | 4.22E-02 |
| chr20 | 62327968 | 62328427 | *RTEL1-TNFRSF6B* | 4 | -4.67 | 0.86 | 6.83E-08 | 3.50E-02 |
| chr4 | 142053146 | 142054660 | *RNF150* | 9 | 114.87 | 20.95 | 4.18E-08 | 2.14E-02 |
| chr20 | 62738880 | 62739073 | *NPBWR2* | 3 | 46.91 | 8.36 | 1.99E-08 | 1.02E-02 |
| chr19 | 55677716 | 55678066 | *DNAAF3* | 7 | 222.67 | 40.89 | 5.15E-08 | 2.64E-02 |
| chr16 | 3493423 | 3493681 | *ZNF597/NAA60* | 6 | -22.95 | 4.20 | 4.71E-08 | 2.41E-02 |
| chr4 | 1161597 | 1161653 | *SPON2* | 2 | 44.62 | 8.17 | 4.75E-08 | 2.43E-02 |
| chr1 | 202113592 | 202114058 | *ARL8A* | 4 | 167.70 | 30.75 | 4.92E-08 | 2.52E-02 |
| chr5 | 140071110 | 140071347 | *HARS2/HARS1* | 5 | 285.97 | 53.45 | 8.81E-08 | 4.51E-02 |
| chr5 | 140529627 | 140530467 | *PCDHB6* | 7 | 15.52 | 2.88 | 6.99E-08 | 3.58E-02 |
| chr14 | 74416980 | 74417249 | *FAM161B/COQ6* | 6 | 266.93 | 49.81 | 8.36E-08 | 4.28E-02 |

Each model adjusted for child sex and age at appetitive trait assessment, batch and estimated white blood cell proportions, maternal age at delivery, smoking during pregnancy, education and pre-pregnancy BMI.
*n* CpGs: number of CpGs included in region; Estimate: effect estimate from meta-analysis of linear regression models.

**Table 3.** Associated regions for epigenome-wide association study of DNA methylation (of cord blood at birth) and emotional undereating at 4-5 years.

| Chromosome | Start (hg19) | End (hg19) | Nearest gene | *n* CpGs | Estimate | SE | *p* | *p*<br>Bonferroni adjusted |
| --- | --- | --- | --- | --- | --- | --- | --- | --- |
| chr4 | 74847646 | 74847829 | *PF4* | 7 | -9.31 | 1.41 | 4.02E-11 | 1.92E-05 |
| chr11 | 368440 | 368638 | *B4GALNT4* | 7 | -29.71 | 4.32 | 5.92E-12 | 2.83E-06 |
| chr13 | 36871878 | 36872346 | *CCDC169-SOHLH2* | 12 | 61.38 | 8.94 | 6.76E-12 | 3.23E-06 |
| chr14 | 100141423 | 100142298 | *HHIPL1* | 5 | -35.00 | 5.60 | 3.94E-10 | 1.89E-04 |
| chr7 | 94286304 | 94286834 | *PEG10* | 19 | 30.63 | 5.18 | 3.42E-09 | 1.64E-03 |
| chr1 | 234367145 | 234367586 | *SLC35F3* | 5 | -11.97 | 2.18 | 3.75E-08 | 1.79E-02 |
| chr2 | 129659420 | 129659834 | *HS6ST1* | 4 | -22.18 | 4.16 | 9.45E-08 | 4.52E-02 |

Each model adjusted for child sex and age at appetitive trait assessment, batch and estimated white blood cell proportions, maternal age at delivery, smoking during pregnancy, education and pre-pregnancy BMI.
*n* CpGs: number of CpGs included in region; Estimate: effect estimate from meta-analysis of linear regression models.

**Table 4.**
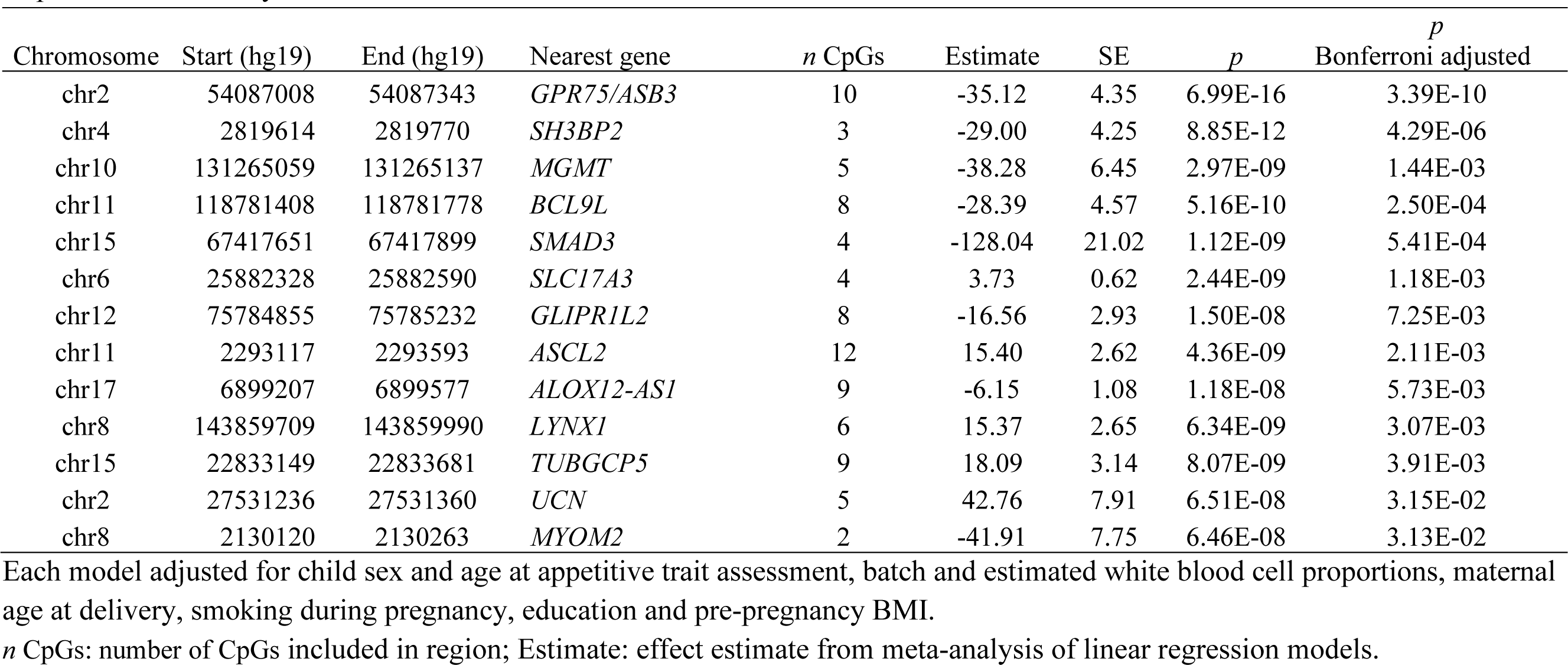
Associated regions for epigenome-wide association study of DNA methylation (of cord blood at birth) and satiety responsiveness at 4-5 years.

**Table 5.** Associated regions for epigenome-wide association study of DNA methylation (of cord blood at birth) and food fussiness at 4-5 years.

| Chromosome | Start (hg19) | End (hg19) | Nearest gene | *n* CpGs | Estimate | SE | *p* | *p*<br>Bonferroni adjusted |
| --- | --- | --- | --- | --- | --- | --- | --- | --- |
| chr1 | 75198582 | 75199117 | *CRYZ/TYW3* | 9 | -18.20 | 2.78 | 5.86E-11 | 2.82E-05 |
| chr6 | 28058724 | 28059208 | *ZSCAN16-AS1* | 9 | -9.14 | 1.33 | 6.88E-12 | 3.31E-06 |
| chr12 | 75784617 | 75785295 | *CAPS2/GLIPR1L2* | 10 | -15.55 | 2.41 | 1.07E-10 | 5.14E-05 |
| chr19 | 57352014 | 57352185 | *ZIM2/MIMT1* | 8 | 72.76 | 10.88 | 2.31E-11 | 1.11E-05 |
| chr4 | 2819614 | 2819770 | *SH3BP2* | 3 | -25.79 | 4.20 | 7.89E-10 | 3.80E-04 |
| chr12 | 2943902 | 2944493 | *NRIP2* | 8 | 14.58 | 2.57 | 1.47E-08 | 7.05E-03 |
| chr2 | 129659018 | 129659946 | *HS6ST1* | 7 | -19.46 | 3.27 | 2.70E-09 | 1.30E-03 |
| chr7 | 150755629 | 150756491 | *SLC4A2* | 10 | -45.95 | 8.49 | 6.16E-08 | 2.96E-02 |
| chr15 | 62516282 | 62516670 | *C2CD4B* | 5 | -15.01 | 2.78 | 6.95E-08 | 3.35E-02 |

Each model adjusted for child sex and age at appetitive trait assessment, batch and estimated white blood cell proportions, maternal age at delivery, smoking during pregnancy, education and pre-pregnancy BMI.
*n* CpGs: number of CpGs included in region; Estimate: effect estimate from meta-analysis of linear regression models.

### 3.2. Enrichment analyses

DNAm associations for emotional undereating, satiety responsiveness, and food fussiness showed some shared regions in the regional-level EWAS, thus confirming the conceptual links between these food avoidant traits. As such, these three food avoidant traits were grouped together for the following enrichment analyses. Hence the following enrichment analyses describe results for ‘food avoidant traits’ and ‘food responsiveness’, the only food approach trait we examined.

#### 3.2.1. Genetic enrichment

Results from the genetic enrichment analyses, which show the extent to which DNAm at regions of associated CpGs might be influenced by genomic variation, are provided in **Supplemental Table 5**. First, we examined whether DNAm at CpGs in each region was related to genetic variation at SNPs (mQTLs, methylation quantitative trait loci). Based on the study by Gaunt et al. (2016), CpGs in regions associated with food avoidant traits were more often related to mQTLs than CpGs in other regions (37.7% *vs.* 7.4%, *p*<0.001), but no difference was found for food responsiveness (14.6% *vs.* 6.9%, *p*=0.08). Based on the study by Min et al. (2020), CpGs in regions associated with both food responsiveness and food avoidant traits were more often related to mQTLs (food responsiveness: 58.7% *vs.* 40.8%, *p*=0.002; food avoidant traits: 81.7% *vs.* 41.6%, *p*<0.001). Second, we examined twin heritability estimates for DNAm of CpGs in each region. The proportions of additive genetic effects estimated were higher for CpGs in associated regions versus those in non-associated regions for both food responsiveness and food avoidant traits (food responsiveness: 0.28 *vs.* 0.14, *p*<0.001; food avoidant traits: 0.52 *vs.* 0.16, *p*<0.001). The proportion of shared environmental effects was smaller for CpGs in associated regions than in other regions for food avoidant traits (0.10 *vs.* 0.16, *p*<0.001), but for food responsiveness no difference was found (0.19 *vs.* 0.19, *p*=0.961). The proportion of unique environmental effects was smaller for CpGs in associated regions than in other regions for both food responsiveness and food avoidant traits (food responsiveness: 0.53 *vs*. 0.67, *p*<0.001; food avoidant traits: 0.38 *vs.* 0.68, *p*<0.001). This means that DNAm at these regions showed evidence of greater influence from genetic rather than environmental variation.

#### 3.2.2. Enrichment for eQTMs

Results from the eQTM look-up, which shows the extent to which CpGs at associated regions have been associated with expression levels of nearby genes in peripheral blood of children (eQTMs), are provided in **Supplemental Table 6**. CpGs at regions associated with food avoidant traits were more often marked as eQTMs than CpGs at other regions (46.8% vs. 4.5%, *p*<0.001). No difference was detected for food responsiveness (10.4% *vs.* 4.7%, *p*=0.08).

#### 3.2.3. Enrichment for regulatory elements

For CpGs in regions associated with food responsiveness, evidence for enrichment of DNAse I hypersensitive regions was found in blood tissue, fetal muscle tissue, fetal stomach tissue, fetal thymus tissue and in induced pluripotent stem cells (*q*<0.05, **Supplemental Figure 1**). No evidence of tissue- or cell-type specific enrichment was found for chromatin states or histone marks for food responsiveness. For food avoidant traits, evidence for enrichment of histone marks was found in the small and large fetal intestine, fetal stomach, fetal trunk muscle, fetal thymus, and fetal adrenal gland (*q*<0.05, **Supplemental Figure 2**). No tissue- or cell-type specific enrichment was found for DNAse I hypersensitive regions or chromatin states with food avoidance traits.

#### 3.2.4. Functional enrichment

Genes associated with differentially methylated regions were tested for functional enrichment against genes associated with all other regions. No enrichment of Gene Ontology pathways was found for any of the appetitive traits (FWER *p*>0.05).

## 4. Discussion

This is the first epigenome-wide association study (EWAS) examining associations of DNA methylation (DNAm) in cord blood with child appetitive traits (age 4 to 5 years), leveraging data from two prospective birth cohorts. The meta-analyses showed that DNAm at individual CpG sites was not associated with any of the four appetitive traits examined in the current study: food responsiveness, emotional undereating, satiety responsiveness and food fussiness. However, multiple differentially methylated regions – genomic regions comprised of several, related methylated sites – were associated with appetitive traits in childhood. Many of the CpG sites in the associated regions have been shown be under greater influence of genetic rather than environmental variation. We also found some evidence that sites associated with appetitive traits were enriched for regulatory elements in tissues such as fetal stomach and intestine. This may indicate a potential regulatory role for the identified differentially methylated regions, but this requires further functional studies. Altogether, these findings provide initial evidence for potential biological pathways underlying the expression of early appetitive traits.

The meta-analyses showed that food responsiveness was associated with the largest number of differentially methylated regions, relative to the other three appetitive traits investigated. This suggests that the prenatal period is a particularly sensitive period for the epigenetic influence of food responsiveness, which may, in part, be due to this construct broadly capturing rudimentary energy balance behaviors which are salient to survival. In children, heightened sensitivity to external food cues observed in food responsiveness serves to promote a surplus in energy intake (Carnell & Wardle, 2007) which favors weight gain (Kininmonth et al., 2021). While adaptive in environments with unreliable energy and nutritional sources, this function may be no longer beneficial in the ubiquitous Western food-environment, where palatable and energy-dense foods are readily available. In our analyses, genes associated with regions differentially methylated for food responsiveness appear to be linked to a broad range of functions, including immune function, neural development, and cardiovascular functioning. In the look-up of genes associated with differentially methylated regions, several notable genes appear to be involved in metabolically-related functions which may be relevant to the early expression of food responsiveness. One example is *ALG9* (ALG9 Alpha-1,2-Mannosyltransferase) on chromosome 11, a gene encoding the glycosylation protein alpha-1,2-mannosyltransferase, which has been shown to be differentially expressed in the placentae of women with diabetes mellitus during pregnancy (Alexander et al., 2018). Another example is a region of chromosome 20, located in the transcription start site of *NPBWR2* (Neuropeptides B/W Receptor 2), which encodes a neuropeptide receptor, that has also been related to leptin and insulin levels in rats (Rucinski et al., 2007) as well as feeding behavior under stress in mice (Aikawa et al., 2008). As food responsiveness was the only food approach trait examined in the current analyses, further research is required to examine associations between DNA methylation and other food approach behaviors, such as enjoyment of food and emotional overeating.

Differentially methylated regions associated with the food avoidance behaviors examined (emotional undereating, satiety responsiveness and food fussiness) were linked to genes also serving broad functions, with a prominence for cardiovascular and gastrointestinal function. Interestingly, several genes were annotated to differentially methylated regions associated with multiple food avoidance behaviors. For example, satiety responsiveness and food fussiness were related to DNAm in or near *SH3BP2* (SH3 Domain Binding Protein 2; chromosome 4) and *GLIPR1L2* (GLIPR1-Like Protein 2; chromosome 12). *SH3BP2* is associated with cherubism, a disorder characterized by dysplasia of the jaw (Reichenberger et al., 2012). *GLIPR1L2* encodes a cysteine-rich secretory protein and is highly expressed in the testes *in vitro*(Ren et al., 2006). Cord blood methylation in *GLIPR1L2* has previously been associated with maternal pre-pregnancy obesity (Martin et al., 2019). In our current analyses, it is worth noting that DNAm in *GLIPR1L2* remained associated with both satiety responsiveness and food fussiness after adjustment for maternal pre-pregnancy BMI (Model 2). This indicates that, where maternal pre-pregnancy BMI may be a potential precursor to cord blood *GLIPR1L2* DNAm, the association between *GLIPR1L2* DNAm and these appetitive traits (satiety responsiveness and food fussiness) is at least partially independent of pre-pregnancy BMI.

Several appetitive traits were associated with DNAm at regions near or at genes linked to pubertal development. Both emotional undereating and food fussiness were associated with DNAm on chromosome 2 at regions near *HS6ST1* (Heparan Sulfate 6-O-Sulfotransferase 1), a gene related to hypogonadism and delayed puberty (Howard et al., 2018). Another differentially methylated region found for emotional undereating was near *IGSF10* (Immunoglobulin Superfamily Member 10) on chromosome 3, which has also been related to delayed puberty (Budny et al., 2020). Additionally, food responsiveness was associated with a differentially methylated region on chromosome 3 in *PLXNB1* (Plexin B1), which has been related to hypogonadism (Welch et al., 2022). Taken together, this could indicate that these gene regions that relate to appetite in early childhood may be signals for growth and development during puberty.

Enrichment analyses showed that differentially methylated regions associated with food avoidance traits were enriched for *cis*-eQTMs in peripheral blood in children, indicating that differential DNAm at these regions may have functional relevance. Among regions in which DNAm was related to expression of the gene closest to the region itself are those at aforementioned *SH3BP2* and *IGSF10*, as well as a region in *OR2L13* (Olfactory Receptor Family 2 Subfamily L Member 13), at which DNAm associated with emotional undereating. This gene encodes an olfactory receptor, and has been shown to be expressed in non-chemosensory tissues as well, including blood and the brain (Ferrer et al., 2016), and DNAm at this gene has been associated with gestational diabetes mellitus previously (Howe et al., 2020; Quilter et al., 2014). DNAm at a few differentially methylated regions was related to expression levels of a gene close to, but not nearest to the region, for example DNAm at *SLC4A2* (Solute Carrier Family 4 Member 2) was associated with food fussiness, yet DNAm at this region is related to the expression of *AGAP3* (ArfGAP With GTPase Domain, Ankyrin Repeat And PH Domain 3), a gene related to synaptic plasticity (Oku & Huganir, 2013).

Further, enrichment analyses of CpGs at differentially methylated regions associated with appetitive traits indicated that, overall, DNAm at the associated regions appeared to be mainly explained by genetic (as opposed to environmental) variation as compared to DNAm at other regions. This corroborates our finding that associations of DNAm with appetitive traits are independent of prenatal factors. Causal pathways could be tested through Mendelian randomization and mediation analyses (at the site-level) to interpret to what extent DNAm might mediate genetic or environmental effects on appetitive traits. Mediation analyses would also be required to formally test appetitive traits as behaviorally-mediated mechanisms linking genetic risk and environmental exposure to weight gain. We did not examine child BMI as an outcome or correlate of appetitive traits, as such analyses are outside the scope of the current analyses as it would require much larger sample sizes. However, this is an important direction for future research, owing to the link between appetitive traits and child BMI and adiposity.

This study adds to the knowledge on biological processes underlying children’s appetitive traits. However, interpretation of these results must be considered in light of some limitations. Firstly, because associations between DNAm at CpG sites and child behavior are typically small (Mulder et al., 2020), we may have been underpowered to detect small effect sizes. However, previous studies examining DNAm in relation to child appetitive traits were small candidate studies (Do et al., 2019; Gardner et al., 2015), and therefore this is the first (and relatively large) EWAS meta-analysis. We encourage researchers to add to these data and to further explore other appetitive traits, as only appetitive traits that were common across both cohorts were examined in the current study. Secondly, appetitive traits were assessed using parent-report, which may introduce subjectivity or social desirability bias. However, the CEBQ was used to assess child appetitive traits, and subscales from the CEBQ have been validated with observations and experimental eating behavior tasks in preschool-aged children (Blissett et al., 2019; Carnell & Wardle, 2007). Parents are likely to be reliable reporters of their child’s behavior as they observe their child’s eating across contexts and over a period of time. Thirdly, there was a 4- to 5- year period between the measurement of the exposure (cord blood DNAm) and the outcome (appetitive traits) in the two cohorts. Other early environmental exposures may influence the expression of children’s eating phenotypes, such as parent feeding practices (Harris et al., 2020). Finally, we only examined one time point of DNAm and appetitive traits. As both appetitive traits and DNA methylation are known to change over time (Derks et al., 2019; Mulder et al., 2021), future research could build on the current findings by examining DNAm and appetitive traits at multiple time points across child development to determine the directionality of associations. There were also many study strengths in addition to those already mentioned. For example, in addition to the analysis of CpGs, we completed a robust analysis of differentially methylated regions. We also controlled for a number of maternal and child covariates in different models, and hypothesized mediators, to examine effect size changes between models.

Findings from this study support the hypothesis that DNAm at numerous genetic regions in cord blood is associated with appetitive traits at preschool-age, implicating widespread DNAm patterns in the newborn as a potential mechanism underlying early childhood eating behavior. We have linked our results to evidence indicating that DNAm at these regions is related to both genetic factors and the prenatal environment, although more so to the former. We hope these findings incite other researchers to study associations between DNAm and appetitive traits, to unravel potential causal pathways through which DNAm may play its role in appetitive traits.

## Acknowledgements

*Generation R*

The Generation R Study is conducted by Erasmus MC, University Medical Center Rotterdam in close collaboration with the School of Law and Faculty of Social Sciences of the Erasmus University Rotterdam, the Municipal Health Service Rotterdam area, Rotterdam, the Rotterdam Homecare Foundation, Rotterdam and the Stichting Trombosedienst & Artsenlaboratorium Rijnmond (STAR-MDC), Rotterdam. We gratefully acknowledge the contribution of children and parents, general practitioners, hospitals, midwives and pharmacies in Rotterdam. The generation and management of the Illumina 450K methylation array data (EWAS data) for the Generation R Study was executed by the Human Genotyping Facility of the Genetic Laboratory of the Department of Internal Medicine, Erasmus MC, the Netherlands. We thank Mr. Michael Verbiest, Ms. Mila Jhamai, Ms. Sarah Higgins, Mr. Marijn Verkerk and Dr. Lisette Stolk for their help in creating the EWAS database. We thank Dr. A.Teumer for his work on the quality control and normalization scripts.

## Author contributions

HAH, PWJ, JFF, and RHM contributed to the conceptualization of the analyses, HAH, CF, AS, JFF, and RHM performed the analyses, HAH, JFF, and RHM interpreted the results and HAH drafted the initial manuscript. All authors reviewed and revised the manuscript, approved the final manuscript as submitted and agree to be accountable for all aspects of the work.

## Declarations of interest

none

## Funding

### Generation R

The general design of Generation R Study is made possible by financial support from Erasmus MC, University Medical Center Rotterdam and the Erasmus University Rotterdam, the Netherlands Organization for Health Research and Development (ZonMW), the Netherlands Organization for Scientific Research (NWO), the Ministry of Health, Welfare and Sport and the Ministry of Youth and Families. The EWAS data were funded by a grant from the Netherlands Genomics Initiative (NGI)/Netherlands Organisation for Scientific Research (NWO) Netherlands Consortium for Healthy Aging (NCHA; project nr. 050-060-810), by funds from the Genetic Laboratory of the Department of Internal Medicine, Erasmus MC, and by a grant from the National Institute of Child and Human Development (R01HD068437). This project received funding from the European Joint Programming Initiative “A Healthy Diet for a Healthy Life” (JPI HDHL, NutriPROGRAM project, ZonMw the Netherlands no.529051022). HAH received funding from the European Union’s Horizon 2020 research and innovation programme under the Marie Skłodowska-Curie grant agreement (No. 707404). The opinions expressed in this document reflect only the authors’ view. The European Commission is not responsible for any use that may be made of the information it contains. PWJ received funding from the Netherlands Organization for Health Research and Development (Mental Health Care Research Program, Fellowship 636320005). The work of RHM and JFF is supported by the European Union’s Horizon 2020 Research and Innovation Programme (EarlyCause; grant agreement No 848158).

### Healthy Start

The Healthy Start study was funded by the National Institute of Diabetes and Digestive and Kidney Diseases (R01DK076648), the National Institute of Environmental Health Sciences (R01ES022934) and the National Institutes of Health Office of the Director (UH3OD023248).

## Data and code availability

For Generation R, data described in the manuscript, code book, and analytic code can be made available upon request to and will be discussed in the Generation R Study Management Team. For Healthy Start, data are available upon reasonable request from the Lifecourse Epidemiology of Adiposity and Diabetes (LEAD) Center data management team at.

## List of abbreviations

CEBQ: Children’s Eating Behaviour Questionnaire
DNAm: DNA methylation
EWAS: Epigenome-wide association study

## Notes

### Competing Interest Statement

The authors have declared no competing interest.

