## Supplemental Figures for "An epigenome-wide association study of child appetitive traits and DNA methylation"

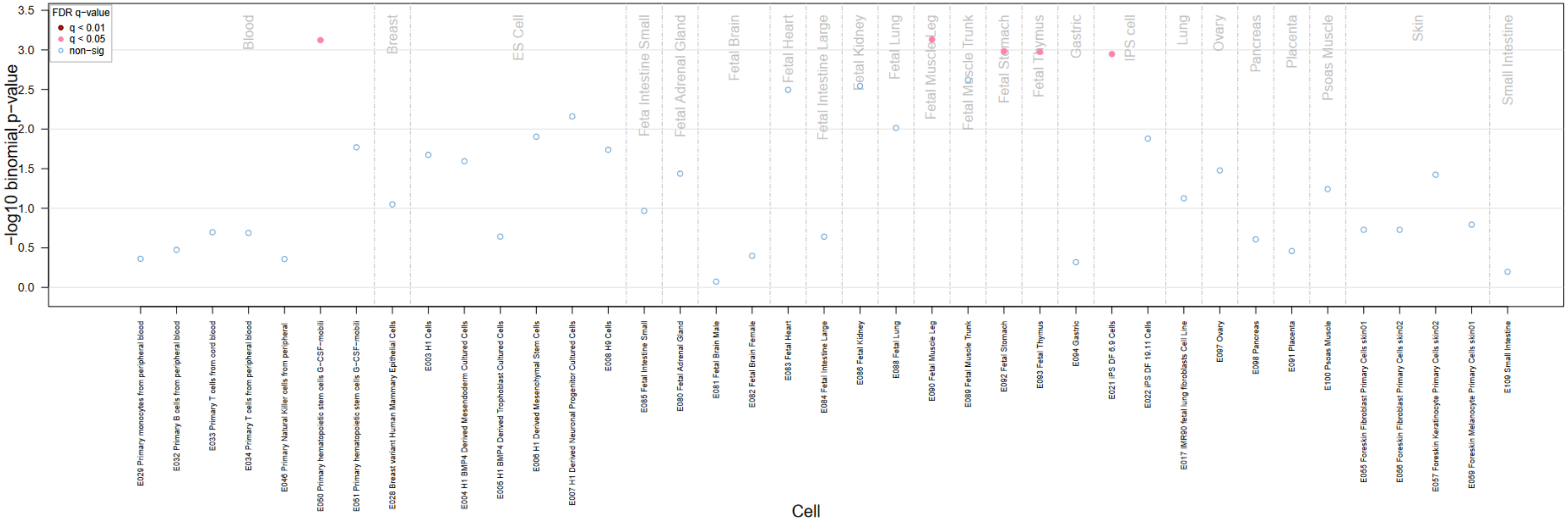


**Supplemental Figure 1.** Enrichment of DNAse I hypersensitive regions co-localization for CpGs in regions associated with food responsiveness

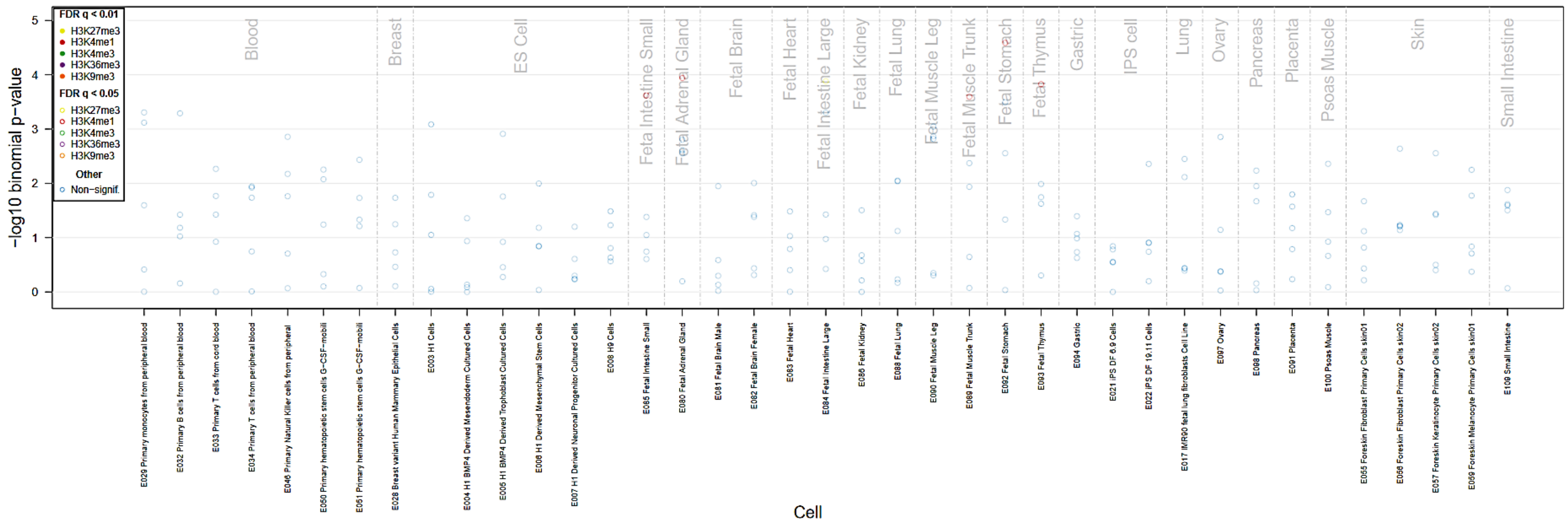


**Supplemental Figure 2.** Enrichment of H3 mark co-localization for CpGs in regions associated with food avoidant traits (emotional undereating, satiety responsiveness, food fussiness)
